# Carbon for nutrient exchange between the lycophyte, *Lycopodiella inundata* and Mucoromycotina ‘fine root endophytes’ is unresponsive to high atmospheric CO_2_ concentration

**DOI:** 10.1101/2020.09.29.318022

**Authors:** Grace A. Hoysted, Jill Kowal, Silvia Pressel, Jeffrey G. Duckett, Martin I. Bidartondo, Katie J. Field

**Author notes:** Corresponding author –.

## Abstract

**Background and Aims:** Non-vascular plants associating with arbuscular mycorrhizal (AMF) and Mucoromycotina ‘fine root endophyte’ (MFRE) fungi derive greater benefits from their fungal associates under higher atmospheric [CO_2_] than ambient, however nothing is known about how changes in [CO_2_] affects MFRE function in vascular plants.

**Methods:** We measured movement of phosphorus (P), nitrogen (N) and carbon (C) between the lycophyte, *Lycopodiella inundata* and Mucoromycotina fine root endophyte fungi using ^33^P-orthophosphate, ^15^N-ammonium chloride and ^14^CO_2_ isotope tracers under ambient and elevated atmospheric [CO_2_] concentrations of 440 and 800 ppm, respectively.

**Key Results:** Transfer of ^33^P and ^15^N from MFRE to plant were unaffected by changes in [CO_2_]. There was a slight increase in C transfer from plant to MFRE under elevated [CO_2_].

**Conclusions:** Our results demonstrate that the exchange of C-for-nutrients between a vascular plant and Mucoromycotina FRE is largely unaffected by changes in atmospheric [CO_2_]. Unravelling the role of MFRE in host plant nutrition and potential C-for-N trade changes between symbionts under varying abiotic conditions is imperative to further our understanding of the past, present and future roles of diverse plant-fungal symbioses in global ecosystems.

## Introduction

Changes in atmospheric CO_2_ concentration ([CO_2_]) have been a prominent feature throughout Earth’s environmental history (Leaky and Lau, 2012). Geochemical models support fossil and stable isotope evidence that the global environment underwent major changes throughout the Palaeozoic Era (541-250 Ma) (Berner et al., 2006; Bergman et al., 2004; Lenton et al., 2016), consisting of a step-wise increase of the Earth’s atmospheric oxygen ([O_2_]), and a simultaneous decline in atmospheric [CO_2_]. Today, Earth faces environmental changes on a similar scale, but with atmospheric [CO_2_] now rising at an unprecedented rate (Meinshausen et al., 2011).

Long before plants migrated onto land, Earth’s terrestrial surfaces were colonised by a diverse array of microbes, including filamentous fungi (Blair, 2009; Berbee et al., 2017). Around 500 Mya, plants made the transition from an aquatic to a terrestrial existence (Morris et al., 2018), facilitated by symbiotic fungi (Nicolson, 1967; Pirosynski and Malloch, 1975). These ancient fungal symbionts are thought to have played an important role in helping the earliest land plants access scarce nutrients from the substrate onto which they had emerged, in much the same way as modern-day mycorrhizal fungi form nutritional mutualisms with plant roots (Pirozynski and Malloch, 1975; Krings et al., 2012). It is highly likely that ancient mycorrhiza-like fungi were closely related to, and subsequently evolved into, modern arbuscular mycorrhizal fungi (AMF) belonging to the fungal subphylum Glomeromycotina (also referred to as phylum Glomeromycota) (Redecker et al.. 2000; Wijayawardene et al., 2018; Radhakrishnan et al., 2020).

Recently, it was discovered that extant non-vascular plants, including the earliest divergent clade of liverworts, associate with a greater diversity of fungi than was previously thought, notably forming endophytic associations with members of the Mucoromycotina (Bidartondo et al., 2011; Desiró et al., 2014; Rimington et al., 2015; Field et al., 2015a). The Mucoromycotina is a partially-saprotrophic fungal lineage (Bidartondo et al., 2011; Field et al., 2015b; 2016) sister to, or pre-dating, the Glomeromycotina AMF, both within Mucoromycota (Spatafora et al., 2017). This discovery, together with the finding that liverwort-Mucoromycotina fungal associations are nutritionally mutualistic (Field et al., 2015a; 2019), often co-occurring with AMF (Field et al., 2016), and emerging fossil evidence (Strullu-Derrien et al., 2014) suggest that the earliest land plants had greater symbiotic options available to them than was previously thought (Field et al., 2015b). Studies now show that symbioses with Mucoromycotina fungi are not limited to early non-vascular plants, but also span almost the entire extant land plant kingdom (Rimington et al., 2015; Orchard et al., 2017a; Hoysted et al., 2018, 2019; Rimington et al., 2020), suggesting that this ancient association may have key roles in modern terrestrial ecosystems.

Latest research into the functional significance of plant-Mucoromycotina fine root endophyte (MFRE) associations indicates that MFRE play a complementary role to AMF by facilitating plant nitrogen (N) assimilation alongside AMF-facilitated plant phosphorus (P) acquisition through co-colonisation of the same plant host (Field et al., 2019). Such functional complementarity is further supported by the observation that MFRE transfer significant amounts of ^15^N but relatively little ^33^P tracers to a host lycophyte *Lycopodiella inundata* in the first experimental demonstration of MFRE nutritional mutualism in a vascular plant (Hoysted et al., 2019). These results contrast with the majority of studies on MFRE and FRE which have, to date, focussed on the role of the fine root endophytes in mediating plant phosphorus (P) acquisition (Orchard et al., 2017b, and literature within).

Today, plant symbiotic fungi play critical roles in ecosystem structure and function. The bidirectional exchange of plant-fixed carbon (C) for fungal-acquired nutrients that is characteristic of most mycorrhizal symbioses (Field and Pressel, 2018) holds huge significance for carbon and nutrient flows and storage across ecosystems (Leake et al., 2004; Rillig, 2004). By forming mutualistic symbioses with the vast majority of plants, including economically-important crops, mycorrhizal fungi have great potential for exploitation within a variety of sustainable management strategies in agriculture, conservation and restoration. Application of diverse mycorrhiza-forming fungi, including both AMF and MFRE, to promote sustainability in agricultural systems is particularly relevant in the context of global climate change and depletion of natural resources (Field et al., 2020). MFRE in particular hold potential for applications to reduce use of chemical fertilisers within sustainable arable systems where routine over-application of N-based mineral fertilisers causes detrimental environmental and down-stream economic impacts (Thirkell et al., 2017). Changes in abiotic factors such as atmospheric [CO_2_] (Cotton, 2018), which is predicted to continue rising in the future (Meinshausen et al., 2011), has been shown to affect the rate and quantities of carbon and nutrients exchanged between mycorrhizal partners (Field et al., 2012, 2015a, 2016; Zheng et al., 2015; Thirkell et al., 2019). As such, insights into the impact of environmental factors relevant to future climate change on carbon for nutrient exchange between symbiotic fungi and plants must be a critical future research goal.

The direct quantification of exchange of plant C and fungal-acquired mineral nutrients, using isotope tracers, has shed important new light on the impact of varying atmospheric [CO_2_] on C-for-nutrient exchanges between plants and their symbiotic fungi (Field et al., 2012, 2015a, 2016; Zheng et al., 2015; Thirkell et al., 2019). Experiments with liverworts associating with MFRE fungi, either in exclusive or in dual symbioses alongside AMF, suggest that these plants derive less benefit in terms of nutrient assimilation from their fungal associates under a high atmospheric [CO_2_] (1,500 ppm) than under a lower [CO_2_] (440 ppm) (Field et al., 2015a; 2016), with the opposite being the case for liverworts associated only with AMF (Field et al., 2012). However, when vascular plant-AMF associations were exposed to high [CO_2_] in the same experimental conditions, there were no changes in mycorrhizal-acquired plant P assimilation (Field et al., 2012). Whether vascular plant-MFRE symbioses respond to changing atmospheric [CO_2_] is completely unknown.

Here, using stable and radioisotope tracers, we investigate how MFRE function in *Lycopodiella inundata*, a homosporous perennial lycophyte widely distributed in the northern hemisphere (Rasmussen and Lawesson, 2002) that associates only with MFRE fungal partners, regarding climate change-relevant shifts in atmospheric [CO_2_]. Specifically, we test the hypotheses that (a) MFRE acquire greater amounts of plant-fixed C under high atmospheric [CO_2_] of 800 ppm as a result of there being larger amounts of photosynthate available for transfer because of greater rates of C fixation by the plant via photosynthesis, and (b), increased C allocation from plant to fungus increases transfer and assimilation of ^15^N and ^33^P tracers from MFRE to plants to feed growing plant demand for nutrients to promote growth.

## Methods

### Plant Material and growth conditions

Mature *Lycopodiella inundata* (L.) plants were collected from Thursley National Nature Reserve, Surrey, UK (SU 90081 39754) in June 2017. The *L. inundata* plants were planted directly into pots (90 mm diameter × 85 mm depth) containing a homogeneous mixture of acid-washed silica sand and 5% pot volume compost (No.2; Petersfield) to aid retention properties of the substrate and to provide minimal nutrients. Soil surrounding plant roots was left intact to prevent damage to the roots and to act as a natural inoculum, including symbiotic fungi and associated microorganisms. Pots were weeded regularly to remove other plant species. All *L. inundata* plants used during this study were collected from the wild.

Based on the methods of Field et al., (2012), three windowed cylindrical plastic cores covered in 10 μm nylon mesh were inserted into the substrate within each experimental pot (Figure 1a-d). Two of the cores were filled with the same substrate as the bulk soil within the pots, comprising a homogeneous mixture of acid-washed silica sand and compost (No. 2; Petersfield), together making up 95% of the core volume, native soil gathered from around the roots of wild plants to ensure cores contained the same microbial communities as in the bulk soil (4% core volume), and fine-ground tertiary basalt (1% core volume) to act as fungal bait (Field et al. 2015a). The third core was filled with glass wool to allow below-ground gas sampling throughout the ^14^C-labelling period to monitor soil community respiration. Plants were watered every other day with distilled water with no other application of nutrient solutions. Microcosms shared a common drip-tray within each cabinet through the acclimation period that ensured a common pool of rhizospheric microorganisms in each microcosm.

**Figure 1.**
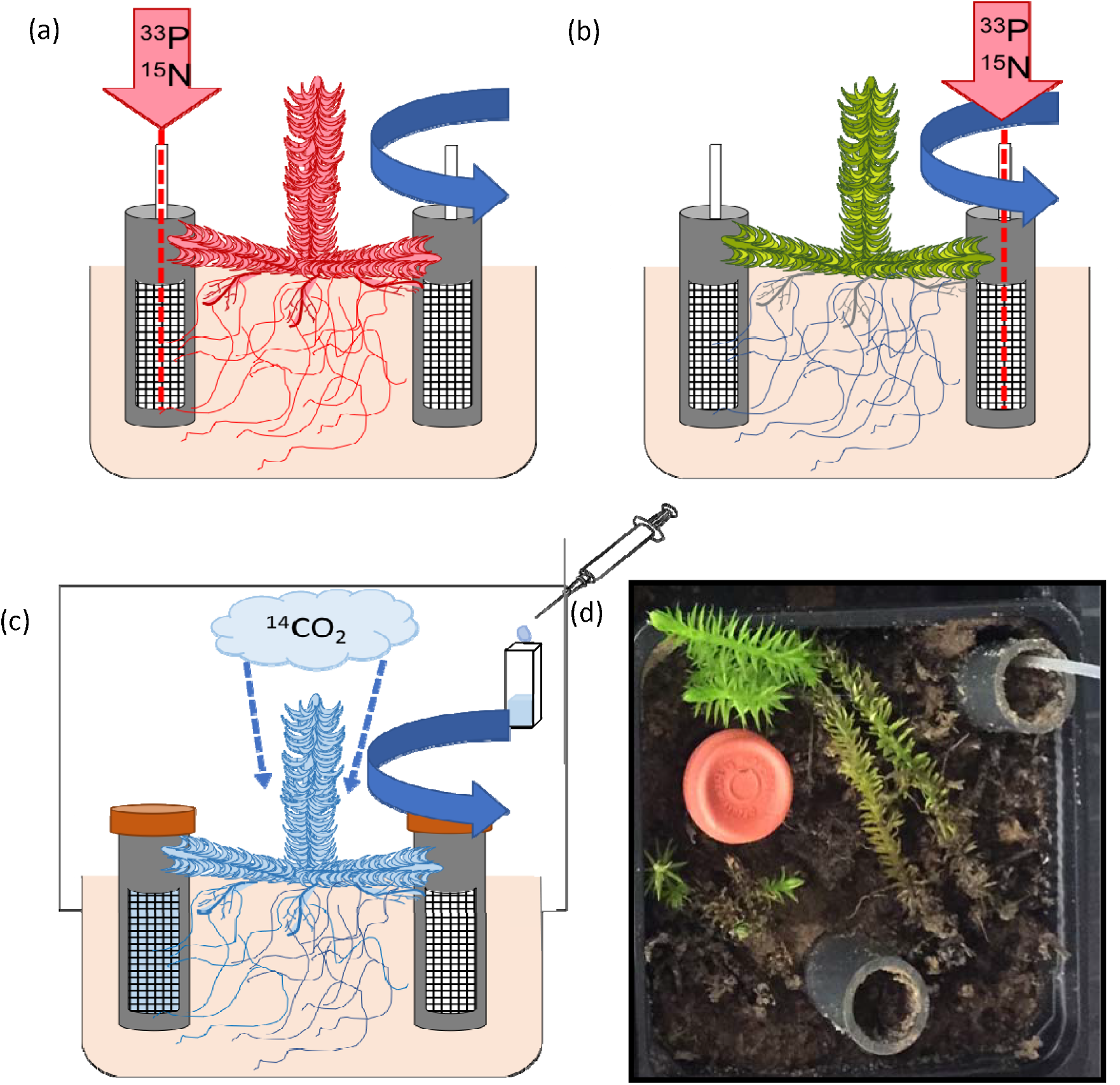
Schematic diagram and photograph of radio and stable isotope tracing system in a lycophyte. (a) Microcosm containing static core in which ^33^P and ^15^N was injected; (b) Microcosm containing rotated core in which ^33^P and ^15^N was injected; (c) Microcosm set-up enclosed in an air-tight container in which ^14^C isotope tracing was conducted and (d) Photograph of *Lycopodiella inundata*-MFRE radio- and stable isotope tracing experimental system.

A total of 48 *L. inundata* microcosms were maintained in controlled environment chambers (Model no. Micro Clima 1200; Snijders Labs) with a light cycle of 16-h daytime (20°C and 70% humidity) and 8-h night-time (at 15°C and 70% humidity). Day-time photosynthetically active radiation (PAR), supplied by LED lighting, was 225 μmol photons m^−2^ s^−1^. Plants were grown at two contrasting CO_2_ atmospheres; 440 ppm [CO_2_] (24 plants) to represent a modern-day atmosphere, or at 800 ppm [CO_2_] (24 plants) to simulate Palaeozoic atmospheric conditions on Earth at the time vascular plants are thought to have diverged (Berner, 2006) as well as predicted atmospheric [CO_2_] for 2100 (Meinshausen et al., 2011). Atmospheric [CO_2_] was monitored using a Vaisala sensor system (Vaisala, Birmingham, UK), maintained through addition of gaseous CO_2_. All pots were rotated within cabinets to control for possible cabinet and block effects. Plants were acclimated to chamber/growth regimes for four weeks to allow establishment of mycelial networks within pots and confirmed by hyphal extraction from soil and staining with trypan blue (Brundrett et al., 1996). Additionally, roots were stained with acidified ink for the presence of fungi, based on the methods of Brundrett et al., (1996). All plants were processed for molecular analyses of fungal symbionts within one week of collection using the protocol in Hoysted et al., (2019).

### Cytological analyses

Roots of experimental *L. inundata* plants were stained with trypan blue (Brundrett et al., 1996), which is common for identifying MFRE (Orchard et al., 2017b), and photographed under a Zeiss Axioscope (Zeiss, Oberkochen, Germany) equipped with a MRc digital camera. To quantify root colonisation by MFRE, five plants were randomly selected per treatment, and from these two intact, healthy roots (per plant) were excised and sectioned transversally in up to six segments (depending on root length) before being processed for SEM according to Duckett et al., (2006). Percentage root colonisation was then calculated by scoring each segment (from a total of 56 and 58 segments respectively for the elevated and ambient [CO_2_] treatments) as colonised or non-colonised under the SEM (Figure 4).

### Quantification of C, ^33^P and ^15^N fluxes between lycophytes and fungi

After the four-week acclimation period, microcosms were moved to individual drip-trays immediately before isotope labelling to avoid cross-contamination of the isotope tracers. A total of 100 μl of an aqueous mixture of ^33^P-labelled orthophosphate (specific activity 111 TBq mmol^−1^, 0.3 ng ^33^P added; Hartmann analytics) and ^15^N-ammonium chloride (1mg ml^−1^; 0.1 mg ^15^N added; Sigma-Aldrich) was introduced into one of the soil-filled mesh cores in each pot through the installed capillary tube. In half (n = 12) of the pots, cores containing isotope tracers were left static to preserve direct hyphal connections with the lycophytes (Fig. 1a). Fungal access to isotope tracers was limited in the remaining half (12) of the pots by rotating isotope tracer-containing cores through 90°, thereby severing the hyphal connections between the plants and core soil (Fig. 1b). These were rotated every second day thereafter, thus providing a control treatment that allows us to distinguish between fungal and microbial contributions to tracer uptake by plants. Assimilation of ^33^P tracer into above-ground plant material was monitored using a hand-held Geiger counter held over the plant material daily.

At detection of peak activity in above-ground plant tissues (21 days after the addition of the ^33^P and ^15^N tracers), the tops of ^33^P and ^15^N-labelled cores were sealed with plastic caps and anhydrous lanolin and the glass wool cores were sealed with rubber septa (SubaSeal, Sigma-Aldrich). Before lights were switched on at 8 am, each pot was sealed into a 3.5 L, gas-tight labelling chamber and 2 ml 10% (w/v) lactic acid was added to 30 μl NaH^14^CO_3_ (specific activity 1.621 GBq/mmol^−1^; Hartmann Analytics), releasing a 1.1-MBq pulse of ^14^CO_2_ gas into the headspace of the labelling chamber (Fig. 1c). Pots were maintained under growth chamber conditions, and 1 ml of headspace gas was sampled after 1 hour and every 1.5 hours thereafter. Below-ground respiration was monitored via gas sampling from within the glass-wool filled core after 1 hour and every 1.5 hours thereafter for ~16 h.

### Plant harvest and sample analyses

Upon detection of maximum below-ground flux of ^14^C, ~16h after the release of the ^14^CO_2_ pulse, each microcosm compartment (i.e. plant material and soil) was separated, freeze-dried, weighed and homogenised. The ^33^P activity in plant and soil samples was quantified by digesting in concentrated H_2_SO_4_ and liquid scintillation (Tricarb 3100TR liquid scintillation analyser, Isotech). The quantity of ^33^P tracer that was transferred to plant by its fungal partner was then calculated using previously published equations (Cameron et al. 2007). To determine total symbiotic fungal-acquired ^33^P transferred to *L. inundata*, the mean ^33^P content of plants that did not have access to the tracer because cores into which the ^33^P was introduced were rotated, was subtracted from the total ^33^P in each plant that did have access to the isotopes within the core via intact fungal hyphal connections (i.e. static cores). This calculation controls for diffusion of isotopes and microbial nutrient cycling in pots, ensuring only ^33^P gained by the plant via intact fungal hyphal connections is accounted for and therefore serves as a conservative measure of the minimum fungal transfer of tracer to the plant.

Between two and four mg of freeze-dried, homogenised plant tissue was weighed into 6 × 4 mm^2^ tin capsules (Sercon) and ^15^N abundance was determined using a continuous flow IRMS (PDZ 2020 IRMS, Sercon). Air was used as the reference standard, and the IRMS detector was regularly calibrated to commercially available reference gases. The ^15^N transferred from fungus to plant was then calculated using equations published previously in Field et al., (2016). In a similar manner as for the ^33^P, the mean of the total ^15^N in plants without access to the isotope because of broken hyphal connections between plant and core contents was subtracted from total ^15^N in each plant with intact hyphal connections to the mesh-covered core to give fungal acquired ^15^N. Again, this provides a conservative measure of ^15^N transfer from fungus to plant as it ensures only ^15^N gained by the plant via intact fungal hyphal connections is accounted for.

The ^14^C activity of plant and soil samples was quantified through sample oxidation (307 Packard Sample Oxidiser, Isotech) followed by liquid scintillation. Total C (^12^C + ^14^C) fixed by the plant and transferred to the fungal network was calculated as a function of the total volume and CO_2_ content of the labelling chamber and the proportion of the supplied ^14^CO_2_ label fixed by plants. The difference in total C between the values obtained for static and rotated core contents in each pot is considered equivalent to the total C transferred from plant to symbiotic fungus within the soil core for that microcosm, noting that a small proportion will be lost through soil microbial respiration. The total C budget for each experimental pot was calculated using equations from Cameron et al., (2006). Total percent allocation of plant-fixed C to extra-radical symbiotic fungal hyphae was calculated by subtracting the activity (in Becquerels (Bq)) of rotated core samples from that detected in static core samples in each pot, dividing this by the sum of activity detected in all components of each microcosm, then multiplying it by 100.

### Statistics

Effects of plant species, [CO_2_] and the interaction between these factors on the C, ^33^P and ^15^N fluxes between plants and Mucoromycotina FRE fungi were tested using analysis of variance (ANOVA) or Mann-Whitney U where indicated. Data were checked for homogeneity and normality. Where assumptions for ANOVA were not met, data were transformed using log_10_. If assumptions for ANOVA were still not met, a Mann-Whitney U as well as a Kruskal-Wallis statistical test was performed. All statistics were carried out using the statistical software package SPSS Version 24 (IBM Analytics).

## Results

### Molecular identification of fungal symbionts

Analysis of experimental *Lycopodiella inundata* plants grown under ambient and elevated [CO_2_] confirmed that they were colonised by Mucoromycotina fine root endophyte fungi. Glomeromycotina fungal sequences were not detected. Mucoromycotina OTUs that have previously been identified in wild-collected lycophytes from diverse locations (Rimington et al., 2015; Hoysted et al., 2019), were detected before and after the experiments. (GenBank/EMBL accession numbers: MK673773-MK673803).

### Cytology of fungal colonisation in plants

Trypan blue staining and SEM of *L. inundata* roots grown under ambient and elevated atmospheric [CO_2_] revealed the same fungal symbiont morphology consistent with that previously observed for *L. inundata*-Mucoromycotina FRE (Hoysted et al., 2019) and MFRE colonisation in other vascular plants (Orchard et al., 2017a) including fine branching, aseptate hyphae (<2 μm diameter) with small intercalary and terminal swellings/vesicles (usually 5-10 but up to 15 μm diameter) but, differently from those in flowering plants, no arbuscules. While no differences were detected in the cytology of colonisation between the roots of plants grown under contrasting atmospheric [CO_2_], those grown under 800 ppm [CO_2_] showed higher percentage colonisation (79%) than those grown under 440 ppm [CO_2_] (53%).

### C transfer from *L. inundata* to MFRE symbionts

The amount of carbon allocated from *L. inundata* to Mucoromycotina FRE fungi under elevated atmospheric [CO_2_] concentrations compared to that when plants were grown under atmospheric [CO_2_] of 440 ppm was greater, with *L. inundata* allocating 2.8 times more recent photosynthate to MFRE fungi in elevated atmospheric [CO_2_] compared with ambient atmospheric [CO_2_], although this was not statistically significant (Figure 2a; Mann-Whitney U = 194, *P* = 0.864, n = 24). In terms of total C transferred from plants to MFRE, a similar trend was observed with *L. inundata* transferring ca. 2.7 times more C to MFRE fungal partners at elevated atmospheric [CO_2_] concentrations of 800 ppm compared to those maintained under atmospheric [CO_2_] of 440 ppm (Figure 2b; Mann-Whitney U = 197.5, *P* = 0.942, n = 24).

**Figure 2.**
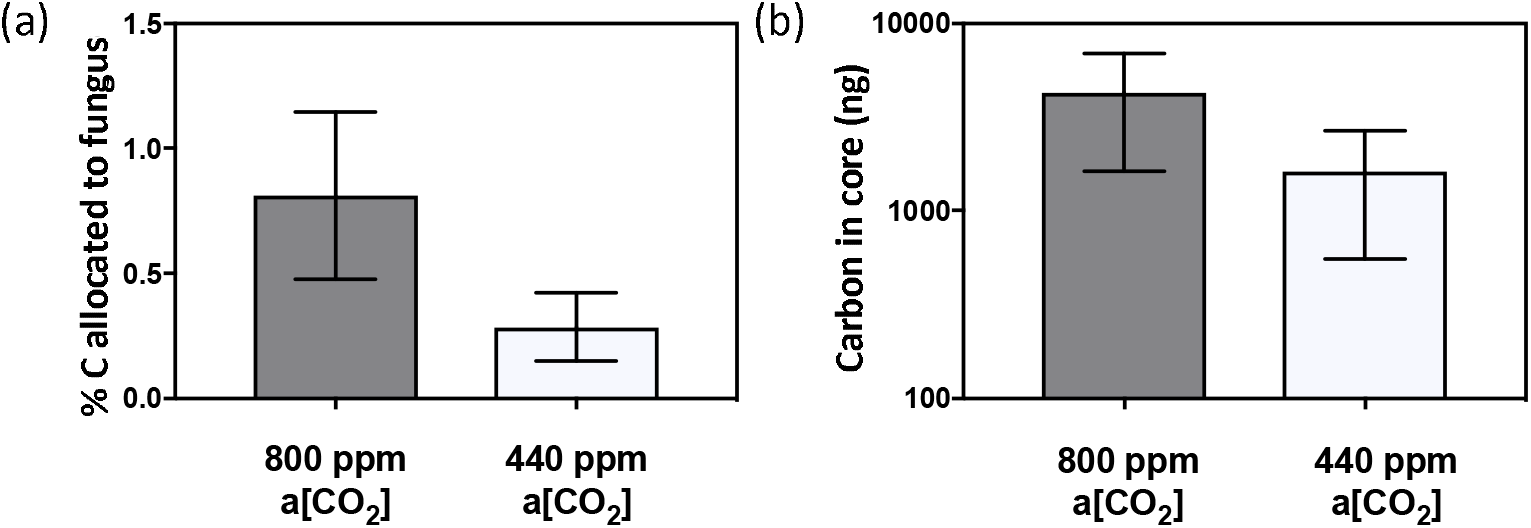
Carbon exchange between *Lycopdiella inundata* and Mucoromycotina fine root endophyte fungi (MFRE). (a) % allocation of plant-fixed C to MFRE; (b) Total plant-fixed C transferred to Mucorocomycotina FRE by *L. inundata*. All experiments conducted at a[CO_2_] of 800 p.p.m. (grey bars) and 440 p.p.m. (white bars). All bars in each panel represent the difference in isotopes between the static and rotated cores inserted into each microcosm. In all panels, error bars denote SEM. In panels (a)-(b) *n* = 24 for both 800 p.p.m and 440 p.p.m a[CO_2_].

### Fungus-to-lycophyte ^33^P and ^15^N transfer

Mucoromycotina FRE transferred both ^33^P and ^15^N to *L. inundata* in both atmospheric CO_2_ treatments (Figure 3). There were no significant differences in the amounts of either ^33^P or ^15^N tracer acquired by MFRE in *L. inundata* plant tissue when grown under elevated [CO_2_] of 800 ppm compared to plants grown under [CO_2_] conditions of 440 ppm, either in terms of absolute quantities (Figure 3a; ANOVA [*F*_1, 23_ = 0.009, *P* = 0.924, N = 10]; Figure 3c; ANOVA [F_1, 22_ = 0.126, *P* = 0.726, n = 10]) or when normalised to plant biomass (Figure 3b; ANOVA [F_1, 23_ = 0.085, *P* = 0.774, n = 10]; Figure 3d; ANOVA [F_1, 22_ = 0.770, *P* = 0.390, n = 10]). Although not significantly different, there was a trend of more nitrogen transferred between MFRE under ambient [CO_2_] compared to elevated [CO_2_] (Figure 3d). Within the experimental microcosms, MFRE transferred 6.65% (± 2.55) ^33^P and 0.07% (±0.02) ^15^N tracer under ambient [CO_2_] and 6.9% (±4.75) ^33^P and 0.03% (±0.18) ^15^N tracer under elevated [CO_2_].

**Figure 3.**
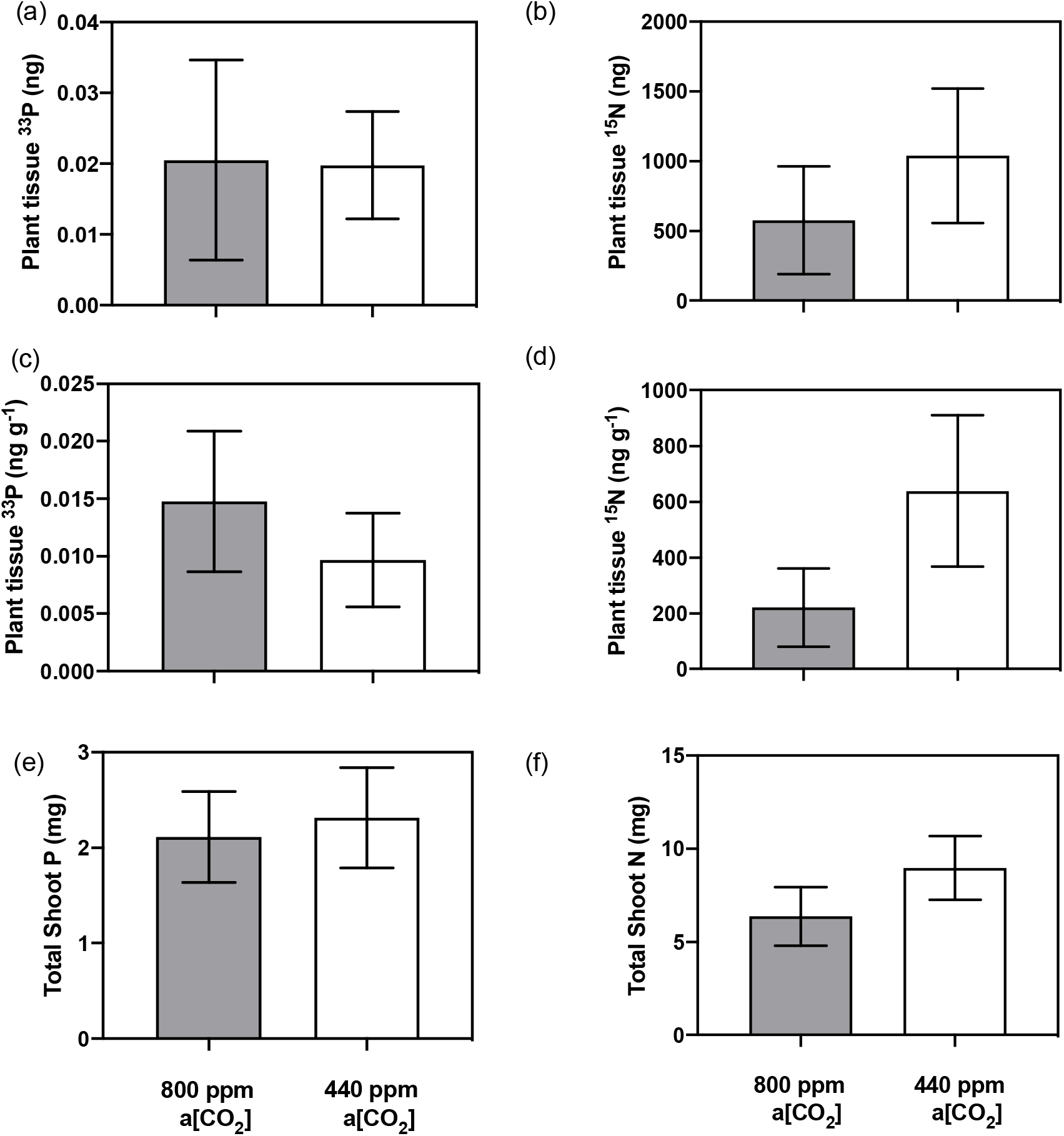
Fungal acquired nutrients by Mucoromycotina fine root endophyte (MFRE) fungi and total shoot nutrients in above ground plant tissue of *L. inundata*. (a) Total plant tissue ^33^P content (ng) and (b) total plant tissue ^15^N content (ng) in *L. inundata* tissue; (c) tissue concentration (ng g^−1^) of fungal-acquired ^33^P and (d) tissue concentration of ^15^N (ng g^−1^) in shoot tissue of *L. inundata*; (e) Total shoot P content (mg) equating to both plant and fungal acquired P and (f) total shoot N content (mg) equating to both plant and fungal acquired N in shoots of *L. inundata*. All experiments conducted at a[CO_2_] of 800 p.p.m. (grey bars) and 440 p.p.m. (white bars). All bars in each panel (a-d) represent the difference in isotopes between the static and rotated cores inserted into each microcosm. In all panels, error bars denote SEM. In panels (a)-(d) *n* = 12 and panels (e)-(f) *n* = 24 for both 800 p.p.m and 440 p.p.m a[CO_2_].

**Figure 4.**
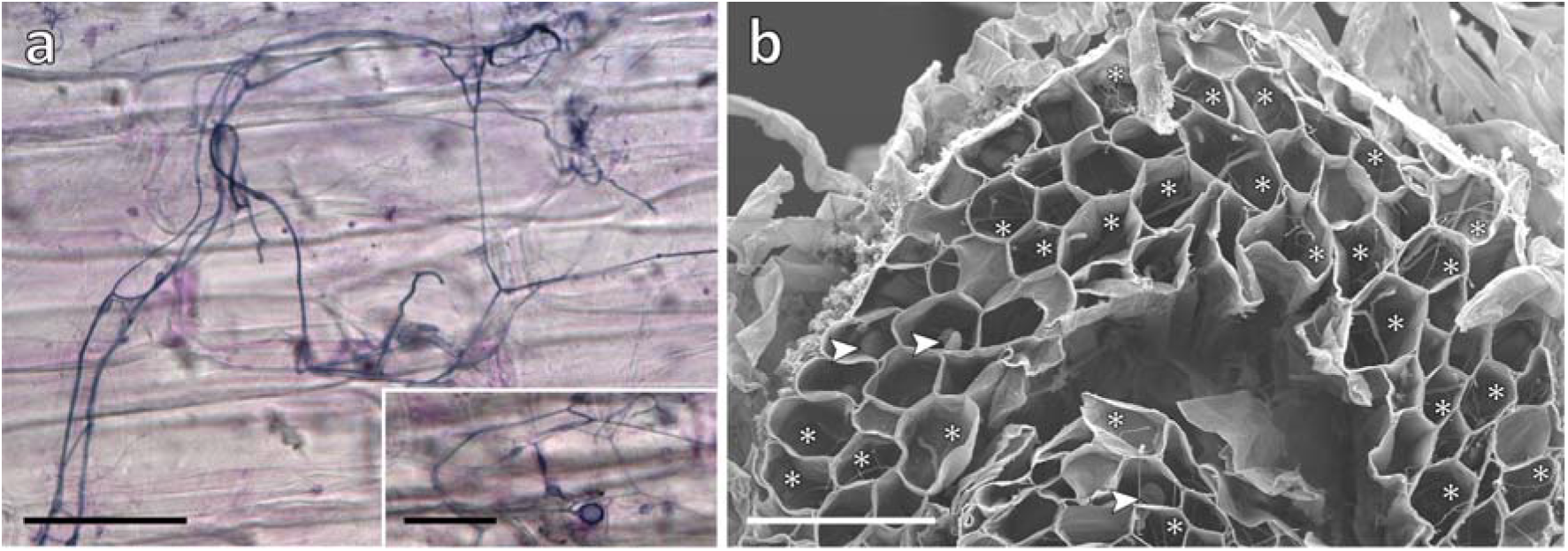
Experimental *Lycopodiella* roots colonised by MFRE. a) Light micrographs of trypan blue stained roots showing fine branching hyphae with intercalary and terminal small vesicles (see insert). b) Scanning electron micrograph of transverse section of root showing abundant branching hyphae (*) and vesicles (arrowed). SEMs at this magnification (x150) were used to quantify % colonisation of roots of experimental plants grown under the two contrasting atmospheric [CO_2_] regimes, shown here in a plant grown under the elevated a[CO_2_] of 800 p.p.m. Scale bars: a) (and (a) insert) 50 μm; b) 100 μm.

## Discussion

Our results demonstrate for the first time, that the exchange of C-for-nutrients between a vascular plant and MFRE symbionts is largely unaffected by changes in atmospheric [CO_2_], with MFRE maintaining ^33^P and ^15^N assimilation and transfer to the plant host across [CO_2_] treatments (Fig. 3a, b), despite MFRE colonisation being more abundant within the roots of plants grown under the elevated [CO_2_] atmosphere. In our experiments, *Lycopodiella inundata* allocated ca. 2.8 times more photosynthate to MFRE under elevated a[CO_2_] compared with plants that were grown under ambient a[CO_2_] (Fig. 2a, b), but without a reciprocal increase in fungal-acquired ^33^P or ^15^N tracer assimilation. Although the difference in C transfer between plants under different [CO_2_] atmospheres was not statistically significant, the trend observed is in line with previous studies where Mucoromycotina fungi (and both Mucoromycotina and AMF partners co-colonising the same host in ‘dual’ symbiosis) fungi gained a greater proportion of recently fixed photosynthates from their non-vascular plant partners, but did not deliver greater amounts of ^33^P or ^15^N tracers when grown under an elevated a[CO_2_] compared to current ambient conditions (Field et al., 2012, 2015a, 2016). This contrasts with patterns of carbon-for-nutrient exchange between other vascular plants and AMF symbionts where increased allocation of carbon to fungal symbionts is usually associated with increases in nutrient delivery from AMF to the host plant (Kiers et al., 2011; Wipf et al., 2019). Such variances may be partly explained by the different lifestyles of the partially saprotrophic Mucoromycotina and the strictly biotrophic Glomeromycotina (Field et al., 2015b, 2016; Field and Pressel, 2018) therefore conventional ‘rules’ governing AMF-plant symbioses may not necessarily apply to Mucoromycotina-plant symbioses. It is possible that when atmospheric [CO_2_] is high, liverworts and relatively simple vascular plants such as *L. inundata* produce photosynthates that they are unable to utilise effectively for growth or reproduction as they possess no or limited vasculature and specialised storage organs to provide transport and storage of excess carbohydrates (Kenrick and Crane, 1997). As a consequence, surplus photosynthates may be either stored as insoluble starch granules, transferred directly to mycobionts (Field et al., 2016), or released into surrounding soil as exudates (Galloway et al., 2017). By moving excess photosynthates into mycobionts, competition for soil resources, space and potential pathogenicity from surrounding saprotrophic organisms such as bacteria and fungi may be reduced (Field et al., 2016).

Additionally, biomass of mycorrhizal fungi may increase in response to elevated a[CO_2_] but this increase does not necessarily result in greater nutrient transfer to the host plant (Alberton et al., 2005), instead inducing a negative feedback through enhanced competition for nutrients between symbiotic partners (Fransson et al., 2007). Studies on ectomycorrhizal fungi, another group of widespread symbiotic fungi with plants that may act as facultative saprotrophs, showed that despite an increase in the amount of extraradical hyphae under elevated [CO_2_], there was no corresponding enhanced transfer of N to the host, suggesting that the fungus had become a larger sink of nutrients (Fransson et al., 2005). While we did not measure fungal biomass in this study, our observation of greater percentage colonisation in the roots of *Lycopodiella* plants grown under an elevated [CO_2_] of 800 ppm compared to those grown under ambient concentrations but with no corresponding increase in fungus-to-plant N and P transfer may suggest a similar possible scenario.

In Field et al. (2015a; 2016), the liverworts *Haplomitrium gibbsiae* and *Treubia lacunosa* transferred approximately 56 and 189 times less photosynthate, respectively, to their MFRE partners under elevated a[CO_2_] than the *L. inundata* in our experiments did – although Field et al.’s (2016) experiments were carried out at higher a[CO_2_] (1,500 ppm) than the present study (800 ppm). Given that *L. inundata* can regulate and maintain CO_2_ assimilation and thus C fixation through stomata and vasculature (Kenrick and Crane, 1997), it is possible that [CO_2_] would affect rates of C fixation and subsequent transfer of plant C to symbiotic fungi less than it might do in poikilohydric liverworts. Our results are consistent with previous studies of AMF symbiosis showing that vascular plants associating with AMF transferred more C to their fungal partners compared to non-vascular species (Field et al., 2012). As such, our findings add to mounting evidence that C transfer from plant to fungus responds differently to changing atmospheric [CO_2_] according to both plant and fungal identity.

We observed no difference in the amount of fungal-acquired ^33^P tracer transferred to *L. inundata* sporophytes between atmospheric [CO_2_] treatments. This aligns with the responses of MFRE symbionts in non-vascular liverworts which also transferred the same amount (or more in the case of *Treubia* mycobionts) of ^33^P and more ^15^N to plant hosts under ambient atmospheric [CO_2_] compared to elevated [CO_2_] (Field et al., 2015a; 2016). The amount of ^33^P transferred to *L. inundata* was much less than has previously being recorded for Mucoromycotina in liverworts (Field et al., 2016) and for Glomeromycotina-associated ferns and angiosperms (Field et al., 2012), despite the same amount of ^33^P being made available, indicating that MFRE fungi do not play a critical role in lycophyte P nutrition. MFRE transferred considerable amounts of ^15^N to their host (see also Hoysted et al., 2019), under both [CO_2_] treatments. This observation together with previous findings that MFRE fungi facilitate the transfer of both organic and inorganic ^15^N to non-vascular liverworts suggest that MFRE may play a complementary role to AMF in plant nutrition, with a more prominent role in N assimilation (Field et al., 2019; Hoysted et al., 2019). While it has been shown that AMF can transfer N to their associated host (Ames et al., 1983; Hodge et al., 2001) significant doubts remain as to the ecological relevance of an AMF-N uptake pathway (see Read 1991; Smith and Smith, 2011). In particular, regarding the exact mechanism of N transfer and, more importantly, the amounts of N transferred via the AMF compared to the N requirements of the plant remain highly equivocal (Smith and Smith, 2011). This murky view of AMF in N transfer, coupled with recent molecular re-identification of fine root endophytes as belonging within Mucoromycotina and not Glomeromycotina (Orchard et al., 2017a) and evidence pointing to a significant role of MFRE in ^15^N transfer to both non-vascular (Field et al., 2015a; 2016; 2019) and vascular plants (Hoysted et al., 2019) suggest that effects in plant N nutrition previously ascribed to AMF may instead be attributed to co-occurring MFRE (Field et al., 2019).

Given that symbioses involving Mucoromycotina FRE fungi are much more widespread than were initially thought, covering a wide variety of habitats (Bidartondo et al., 2011; Rimington et al., 2015, 2020; Orchard et al., 2017a, b, c; Hoysted et al., 2018; 2019), their role in plant N nutrition and responses to high [CO_2_] may have much broader ecological significance than previously assumed. It remains critical that we test how mycorrhizal plasticity (both AMF and FRE) translates into function in order to understand how climate change may affect nutrient fluxes between symbionts in the past, present and, importantly, the future (Field and Pressel, 2018). With the growing interest in the application of mycorrhizal fungi to improve vascular plant community diversity, increasing MFRE in soils alongside AMF may improve host nutrient capture and ecosystem robustness, including conservation of endangered species, such as *L. inundata*, in the face of global climate change (van der Heijden et al., 1998; 2015).

In this study, we provide a first assessment of the effects of varying [CO_2_] on carbon-for-nutrient exchanges between MFRE and a vascular plant. Our results point to important differences in responses to changing [CO_2_] between MFRE and AMF and between MFRE symbioses in vascular vs. non-vascular plants, however they are restricted to one, early diverging, vascular species, generally growing in severely N-limited habitats. It is now critical that similar investigations are extended to a broader range of taxa, including flowering plants known to engage in symbiosis with both MFRE and AMF. In doing so, efforts towards the potential exploitation of these symbiotic fungi to help meet sustainability targets of the future may be better informed and the likelihood of success vastly improved.

## Acknowledgements

We gratefully acknowledge support from the NERC to KJF, SP (NE/N00941X/1; NE/S009663/1) and MIB (NE/N009665/1). KJF is funded by a BBSRC Translational Fellowship (BB/M026825/1) and a Philip Leverhulme Prize (PLP-2017-079). We thank Natural England for granting permission to collect *Lycopodiella* from Thursley.

